# Evolution of cold acclimation in temperate grasses (Pooideae)

**DOI:** 10.1101/210021

**Authors:** Marian Schubert, Lars Grønvold, Simen R. Sandve, Torgeir R. Hvidsten, Siri Fjellheim

## Abstract

In the past 50 million years climate cooling has triggered the expansion of temperate biomes. During this period, many extant plant lineages in temperate biomes evolved from tropical ancestors and adapted to seasonality and cool conditions. Among the Poaceae (grass family), one of the subfamilies that successfully shifted from tropical to temperate biomes is the Pooideae (temperate grasses). Subfamily Pooideae contains the most important crops cultivated in the temperate regions including wheat (*Triticum aestivum*) and barley (*Hordeum vulgare*). Due to the need of well-adapted cultivars, extensive research has produced a large body of knowledge about the mechanisms underlying cold adaptation in cultivated Pooideae species. Especially cold acclimation, a process which increases the frost tolerance during a period of non-freezing cold, plays an important role. Because cold adaptation is largely unexplored in lineages that diverged early in the evolution of the Pooideae, little is known about the evolutionary history of cold acclimation in the Pooideae. Here we test if several species of early diverging lineages exhibit increased frost tolerance after a period of cold acclimation. We further investigate the conservation of five well-studied gene families that are known to be involved in the cold acclimation of Pooideae crop species. Our results indicate that cold acclimation exists in early diverging lineages, but that genes involved in regulation of cold acclimation are not conserved. The investigated gene families show signs of lineage-specific evolution and support the hypothesis that gene family expansion is an important mechanism in adaptive evolution.

## Introduction

Temperate biomes expanded during global cooling throughout the Eocene and the Eocene to Oligocene (E-O) transition, *ca.* 34 million years ago (Mya) (Potts and Behrensmeyer 1992, Donoghue 2008, Stickley et al. 2009, Mudelsee et al. 2014). In plants, biome shifts are rare and restricted to a small number of lineages (Crisp et al. 2009) and only a few, ancestrally tropical angiosperm lineages colonized the expanding temperate biomes (Judd et al. 1994, Wiens and Donoghue 2004, Kerkhoff et al. 2014). The successful colonizers faced several stresses and adapted to frost, increased temperature seasonality, and short growing seasons (Zachos et al. 2001, Eldrett et al. 2009, Mudelsee et al. 2014).

The grass (Poaceae) flora of temperate biomes is dominated by members of the subfamily Pooideae which account for up to 90% of the grass species in Northern biomes (Hartley 1973; Clayton 1981). Since their emergence in the late Paleocene or early Eocene (Bouchenak-Khelladi et al. 2010, Christin et al. 2014, Spriggs et al. 2014), the grass subfamily Pooideae successfully shifted their distribution from the tropical/subtropical biomes of their ancestors (Edwards and Smith 2010; Strömberg 2011) to temperate biomes. Except for a few hundred species in early diverging Pooideae tribes most of the *ca.* 4200 Pooideae species belong to the species-rich ‘core Pooideae’ clade (*sensu* Davis and Soreng 1993; Soreng and Davis 1998) and its sister tribe Brachypodieae, which contains the model grass *Brachypodium distachyon* (Soreng et al. 2015). Because core Pooideae contain economically important species like wheat (*Triticum aestivum*) and barley (*Hordeum vulgare*) as well as several forage grasses like ryegrass (*Lolium perenne*), the need for well-adapted cultivars has provided extensive knowledge about molecular mechanisms underlying adaptation to temperate environments (Thomashow 1999; Sandve et al. 2008; Galiba et al. 2009; Sandve et al. 2011, Preston and Sandve 2013; Fjellheim et al. 2014). Key molecular responses to low temperatures in Pooideae species may represent adaptations to temperate climate (McKeown et al. 2016, Woods et al. 2016, McKeown et al. 2017), whose expansion coincided with the early evolution of the Pooideae.

Plants in temperate biomes respond to frost and periods of prolonged cold with a suite of physiological and biochemical adaptations controlled by complex regulatory programs. The main challenge of exposure to frost is maintaining the integrity of the cellular membranes to avoid dehydration (Pearce et al. 2001). This is accomplished by adjusting the lipid composition of membranes to increase membrane stability (Uemura et al. 1995; Danyluk et al. 1998;) and to accumulate sugars and anti-freeze proteins (Griffith and Yaish 2004; Sandve et al. 2011). Additionally, accumulation of reactive oxygen species (ROS) damages lipid membranes and increases protein degradation (Murata et al. 2007; Crosatti et al. 2013) to which plants react by synthesizing proteins that decrease ROS-mediated stress (Crosatti et al. 2013). Through the process of cold acclimation – a period of cold, non-freezing temperatures – temperate plants can increase their frost tolerance to prepare for freezing during winter (Thomashow 1999; Thomashow 2010).

Five of the best studied cold acclimation gene families code for C-repeat binding factors (CBF), dehydrins (DHN), chloroplast-targeted cold-regulated proteins (ctCOR), ice recrystallization inhibition proteins (IRIP) and fructosyl transferases (FST). All of these are known to be induced by cold and play important roles during cold-stress response and cold acclimation in core Pooideae (*CBF*: Badawi et al. 2007; Li et al. 2012. *DHN*: Olave-Concha et al. 2004; Rorat 2006; Kosová et al. 2007, 2014. *ctCOR*: Gray et al. 1997; Crosatti et al. 1999, 2013; Tsvetanov et al. 2000. *IRIP*: Antikainen and Griffith 1997; Hisano et al. 2004; Kumble et al. 2008; Sandve et al. 2008; John et al. 2009; Zhang et al. 2010; Sandve et al. 2011. *FST*: Hisano et al. 2004; Tamura et al. 2014). The CBFs are transcription factors and function as “master-switches” of cold regulation and cold acclimation (Sarhan et al. 1998; Thomashow 1999) and are involved in various kinds of stress response (Agarwal et al. 2006; Akhtar et al. 2012). Two groups of *CBF* genes – *CBFIII* and *CBFIV* – are especially important for cold acclimation in Pooideae and are restricted to this subfamily (Badawi et al. 2007, Li et al. 2012). The *DHN* gene family encodes hydrophilic proteins that share a lysine-rich sequence, the “K-segment”, which interacts with membranes (Koag et al. 2003; Koag et al. 2009) and protects against dehydration-related cellular stress and possibly also acts as cryoprotectant (Close 1997; Danyluk et al. 1998; Houde et al. 2004). ctCOR proteins (i.e. COR14 and WCS19; Gray et al. 1997; Crosatti et al. 1999; Tsvetanov et al. 2000) are thought to alleviate damage by ROS that accumulate during cold-induced overexcitation of the photosystems (Crosatti et al. 2013). IRIPs bind to the edges of microscopic intracellular ice grains and restrict the formation of large hazardous ice crystals (Griffith and Ewart 1995; Antikainen and Griffith 1997; Sidebottom et al. 2000; Griffith and Yaish 2004; Tremblay et al. 2005; Sandve et al. 2011). Lastly, frost tolerance correlates with the accumulation and degree of polymerization of fructans which are synthesized by FSTs (Hisano et al. 2004; Tamura et al. 2014). Fructans are the major carbohydrate storage in model Pooideae species (Chalmers et al. 2005), but they have also been shown to improve membrane stability during freezing stress (Hincha et al. 2000). Interestingly, Li et al. (2012) identified cold responsive *IRIP* and *CBFIII* homologs in the model grass *B. distachyon*, which is sister to the core Pooideae lineage, while homologous *CBFIV* and *FST* genes were absent.

Because our knowledge about cold adaptations in Pooideae mostly stems from studies restricted to a handful species in the core Pooideae-Brachypodieae clade, it is unknown when and how cold acclimation and responsible genes evolved. A prerequisite for a successful biome shift lies in a lineage’s adaptive potential, which is controlled by the genetic toolkit of its ancestor (Edwards and Donoghue 2013; Christin et al. 2015). Therefore, functional knowledge about the five gene families provides an excellent basis to investigate their importance for the evolution of cold acclimation in the Pooideae. If those genes have been part of the cold response in the most recent common ancestor (MRCA) of the Pooideae, we expect to find a high degree of conservation and same expression patterns among Pooideae species. Alternatively, if cold adaptation evolved independently in the main Pooideae lineages, we expect a lower degree of conservation and a more diverse expression pattern.

Here we test the hypotheses that cold acclimation is conserved in Pooideae and that key cold acclimation genes known from core Pooideae are cold responsive in early diverging lineages. To test the conservation of cold acclimation we performed a classic growth experiment and investigated the effect of cold acclimation across the three core Pooideae species, three Brachypodieae species and three early diverging Pooideae species *Nardus stricta*, *Melica nutans* and *Stipa lagascae*. To test conservation of cold acclimation genes we compared the expression of *CBF*, *DHN*, *ctCOR*, *IRIP* and *FST* homologs between *H. vulgare*, *B. distachyon* and the three early diverging Pooideae species. All species increase their frost tolerance following cold acclimation, but only two out of the ten studied genes exhibited completely conserved expression pattern across the investigated species. Nevertheless, we suggest that gene families *DHN*, *ctCOR*, *CBFIIId* and *CBFIV* were instrumental in the Pooideae’s shift to temperate biomes. The Pooideae MRCA might not have been cold tolerant, but those gene families were likely part of its genetic toolkit that steered the evolutionary trajectory of its descendants towards cold tolerance.

## Material and methods

### Plant material and freezing tests

Seeds from nine Pooideae species were acquired from germplasm collections or collected in nature. From the core Pooideae we included *Hordeum vulgare* L. (cultivar ‘Sonja’, provided by Professor Åsmund Bjørnstad, Department of Plant Sciences, Norwegian University of Life Sciences, Ås NO-1432, Norway), *Lolium perenne* L. (cultivar ‘Fagerlin’, provided by Dr. Kovi, Department of Plant Sciences, Norwegian University of Life Sciences, Ås NO-1432, Norway) and *Elymus repens* (L.) Gould (collected in October 2015, Norway [59.66111, 10.89194]). From the tribe Brachypodieae we included *Brachypodium distachyon* (L.) P. Beauv. (accession ‘Bd1-1’ [W6 46201] acquired from U.S. National Plant Germplasm System [U.S.-NPGS] via Germplasm Resources Information Network [GRIN]), *B. pinnatum* (collected in October 2015, Norway [59.71861, 10.59333]) and *B. sylvaticum* (collected in October 2015, Norway [59.68697, 10.61012]). From early diverging Pooideae lineages we included *Melica nutans* L. (collected in June 2012, Germany [50.70708, 11.23838]), S*tipa lagascae* Roem & Schult. (accession PI 250751, acquired from U.S.-NPGS via GRIN), and *Nardus stricta* (collected in July 2012, Romania [46.69098, 22.58302]).

The seeds were germinated and plants grown in the green house at 20°C under natural day light. Each individual was divided into four clones, one for each treatment and control. The plants were acclimated at 4°C and short (8h) days for three weeks. Control conditions were short days and 20°C. The light intensity was 50 µmol m^−2^ s^−1^. At the end of the cold acclimation period, plants were subjected to freezing at three different temperatures (−4, −8 and −12°C) following Alm et al. (2011). For each temperature we used 15 acclimated and 15 non-acclimated individuals per species. Additional 15 individuals per species were kept at control conditions. After freezing, plants were cut down to approximately 3 cm and grown at 20°C under long days in a greenhouse with natural light conditions. Two and three weeks after the plants were moved into 20°C and long days they were assessed for regeneration ability and scored from 0 (dead) to 9 (growth without damage). Differences between acclimated and non-acclimated individuals within each species were tested with a one-tailed *t* test in R (R Core Team 2016) using the ‘stats’ package.

### Transcriptomic data and differential gene expression

In another study (Grønvold et al., 2017) we compared the molecular cold response from the five Pooideae species *Nardus stricta, Stipa lagascae*, *Melica nutans, Brachypodium distachyon* (same populations/accessions as described above) and *Hordeum vulgare* (cultivar ‘Igri’, provided by Prof. Åsmund Bjørnstad). The transcriptomic data produced in Grønvold et al. 2017 was used as basis for the investigation of cold acclimation genes.

In short, seeds from the five species were germinated and initially grown in greenhouse at a neutral day length (12 hours of light), 17°C and a minimum light intensity of 150 µmol m^−2^ s^−1^. After an initial growth period plants were randomly distributed to two growth chambers and subjected to cold treatment (6°C) under short days (8 hours of light) and a light intensity of 50 µmol m^−2^ s^−1^. Leaf material for RNA isolation was collected after eight hours, four weeks and nine weeks of cold treatment. The sampling points were chosen to separate between short-term and long-term cold responsive expression patterns. For each time point, flash-frozen leaves from five individuals were individually homogenized using a TissueLyser (Qiagen Retsch, Haan, Germany) and total RNA was isolated (from each individual) using RNeasy Plant Mini Kit (Qiagen Inc., Germany), following the manufacturer’s instructions. Purity and integrity of total RNA extracts was determined using a NanoDrop 8000 UV-Vis Spectrophotometer (Thermo Scientific, Wilmington, DE, USA) and 2100 Bioanalyzer (Agilent, Santa Clara, CA, USA), respectively. Pooled RNA extracts were delivered to the Norwegian Sequencing Centre (NSC), Centre for Evolutionary and Ecological Synthesis (CEES), Department of Biology, University of Oslo, Norway, where strand-specific cDNA libraries were prepared and paired end sequenced on a HiSeq 2000-system (Illumina, San Diego, CA, USA). *De novo* transcriptomes were assembled using Trinity v2.0.6 (Grabherr et al. 2011). Ortholog groups were inferred with orthoMCL (Li et al. 2003) using protein sequences translated from the assembled transcriptomes in addition to reference genomes from *H. vulgare*, *Lolium perenne* and *B. distachyon* as well as *Oryza sativa, Sorghum bicolor* and *Zea mays*. For each *de novo* transcriptome differential gene expression was estimated using DESeq2 (Love et al. 2014) for transcripts belonging to an ortholog group. We used an in-house computational pipeline to identify high confidence ortholog groups based on phylogenetic analysis of the orthoMCL ortholog groups.

### Identification of cold acclimation candidate genes

For five cold acclimation gene families (*C-repeat binding factor* genes (*CBFIII* and *CBFIV*), *dehydrin* genes (*DHN*), *chloroplast targeting cold-regulated* genes (*ctCOR*), *ice-recrystallization inhibition protein* genes (*IRIP*) and *fructosyltransferase* genes (*FST*)) we extracted previously identified *H. vulgare* genes (Table S1) and downloaded translated amino acid (AA) from GenBank. These reference sequences were used in protein BLAST searches against translated *de novo* transcript sequences produced by Grønvold et al. 2017. Potential cold acclimation gene transcripts were identified by discarding protein BLAST results with a maximum bitscore < 90, and e-value > 1E-021. When at least one transcript met those criteria, all transcripts of the respective high confidence ortholog group were defined as candidate transcripts. The high confidence ortholog groups also contained genomic coding sequences (CDS) from the Pooideae species *H. vulgare, L. perenne* and *B. distachyon* as well as from the outgroup species *O. sativa*, *S. bicolor* and *Z. mays*. *De novo* transcripts not part of a high confidence ortholog group were included when they met a maximum bitscore > 110 and an e-value < 1E-31. Since seven of the *H. vulgare* transcripts were found that did not belong to any ortholog groups, differential expression had to be re-run for *H. vulgare* after including these transcripts. MUSCLE v.3.8.31 (Edgar 2004) was used to create multiple sequence alignments by aligning all nucleotide candidate transcripts with *H. vulgare* reference coding sequences (CDS) and best CDS nucleotide BLAST hits for *Triticum sp.* acquired from GenBank. To reduce redundancy, genomic *H. vulgare* reference sequences of the ortholog groups were removed from the alignments when they were identical to *H. vulgare* sequences from GenBank.

To ensure proper tree root resolution, further outgroups were included where necessary. Alignments were manually trimmed and optimized using AliView v1.7.1 sequence editor (Larsson 2014). Transcripts were excluded from subsequent analyses when i) cross contamination was identified, i.e. very low total expression and sequence nearly identical to highly expressed contamination source or ii) the transcript was too fragmented/truncated to contain sufficient phylogenetically informative characters.

### Gene tree reconstruction and calibration

For each multiple sequence alignment, the best model of nucleotide substitution (either HKY + Γ, or GTR + Γ) was chosen based on estimations of jModelTest v2.1.7 (Darriba et al. 2012). Using BEAST v1.8.2 (Drummond et al. 2012), we estimated gene trees under an uncorrelated lognormal relaxed clock model with the prior mean substitution rate uniformly distributed between 0 and 0.06 and a Yule tree prior (birth only) for 100 Million MCMC generations while model parameters were logged every 100000 generations. Tracer v1.6 (http://tree.bio.ed.ac.uk/software/tracer/) was used to assure that all parameters had an effective sample size (ESS) above 200. Ten percent of all trees were discarded (burn-in). The remaining trees were concatenated to the maximum clade credibility tree by TreeAnnotator v2.3.0 (Drummond and Rambaut 2012) and visually adjusted using FigTree v1.4.2 (http://tree.bio.ed.ac.uk/software/figtree/). Posterior probabilities equal to or greater than 0.8 were considered to be a significant node support. During BEAST analyses, two normally distributed node age priors were used to calibrate gene trees. Prior for *H. vulgare* and *B. distachyon* divergence was set to 44.4 My (3.53 standard deviation) according to estimates from Marcussen et al. (2014) and priors for *O. sativa* and *B. distachyon* divergence was set to 53 Mya (3.6 standard deviation) according to estimates from Christin et al. (2014). To ensure correct rooting of the gene trees, we constrained Pooideae transcripts to form a monophyletic clade with their closest *O. sativa* sequences.

## Results

### Freezing tests

Freezing tests revealed that cold acclimation, i.e. increased frost tolerance through exposure to cold non-freezing periods, exists in species of early diverging lineages. All acclimated plants from those lineages exhibited a higher survival rate at −4 and −8°C compared to non-acclimated plants (Fig. 1), although the *t* test was not significant at −4°C for *S. lagascae* and *N. stricta*. Interestingly, non-acclimated plants of early diverging Pooideae species performed better at −4°C than non-acclimated *Brachypodium* species and *H. vulgare*, comparable with the survival rates of non-acclimated, perennial core Pooideae species *L. perenne* and *Elymus repens*.

**Figure 1:**
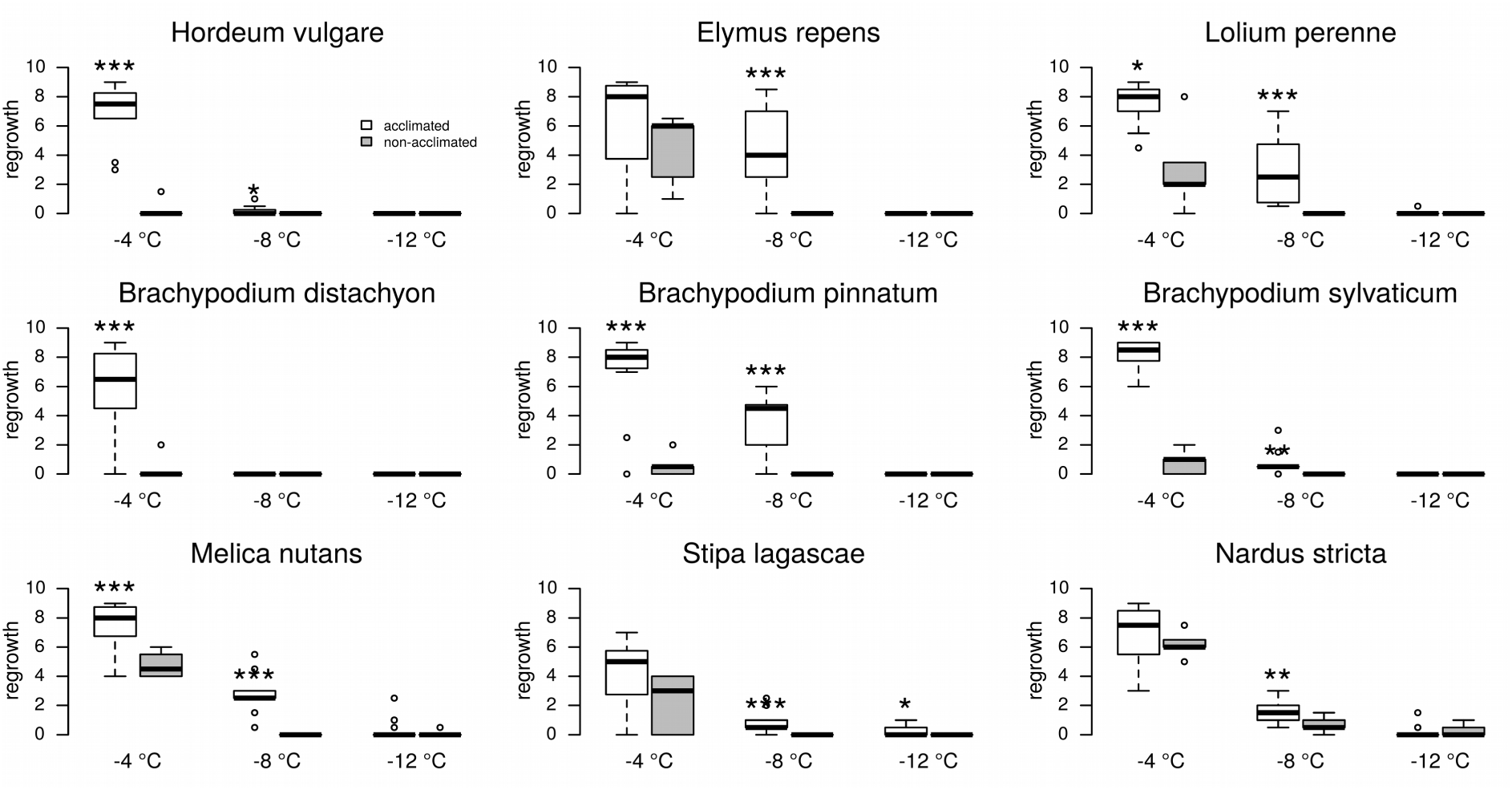
Frost tolerance after cold acclimation. Regrowth of nine acclimated and non-acclimated Pooideae species after exposure to three freezing temperatures (−4°C, −8°C and 12°C). Significant differences in regrowth between acclimated and non-acclimated plants are indicated by asterisks (*** p ≤ 0.001, ** p ≤ 0.01, * p ≤ 0.05).

### *CBFIIIc/d* and *CBFIV* gene family

Consistently with previous results (Badawi et al. 2007), we reconstructed two monophyletic, Pooideae-specific *CBFIII* clades (Fig. 2). In the *CBFIIIc* clade, only *de novo* transcripts from *H. vulgare* were represented but none of them were differentially expressed. The *CBFIIId* clade contained numerous *de novo* transcripts from all studied species, but did not clearly reflect the Pooideae’s species phylogeny. All but three *de novo* transcripts were differentially expressed and either induced in short-term cold (*N. stricta* and H. *vulgare*) or in short- and long-term cold (*B. distachyon* and *M. nutans*).

**Figure 2:**
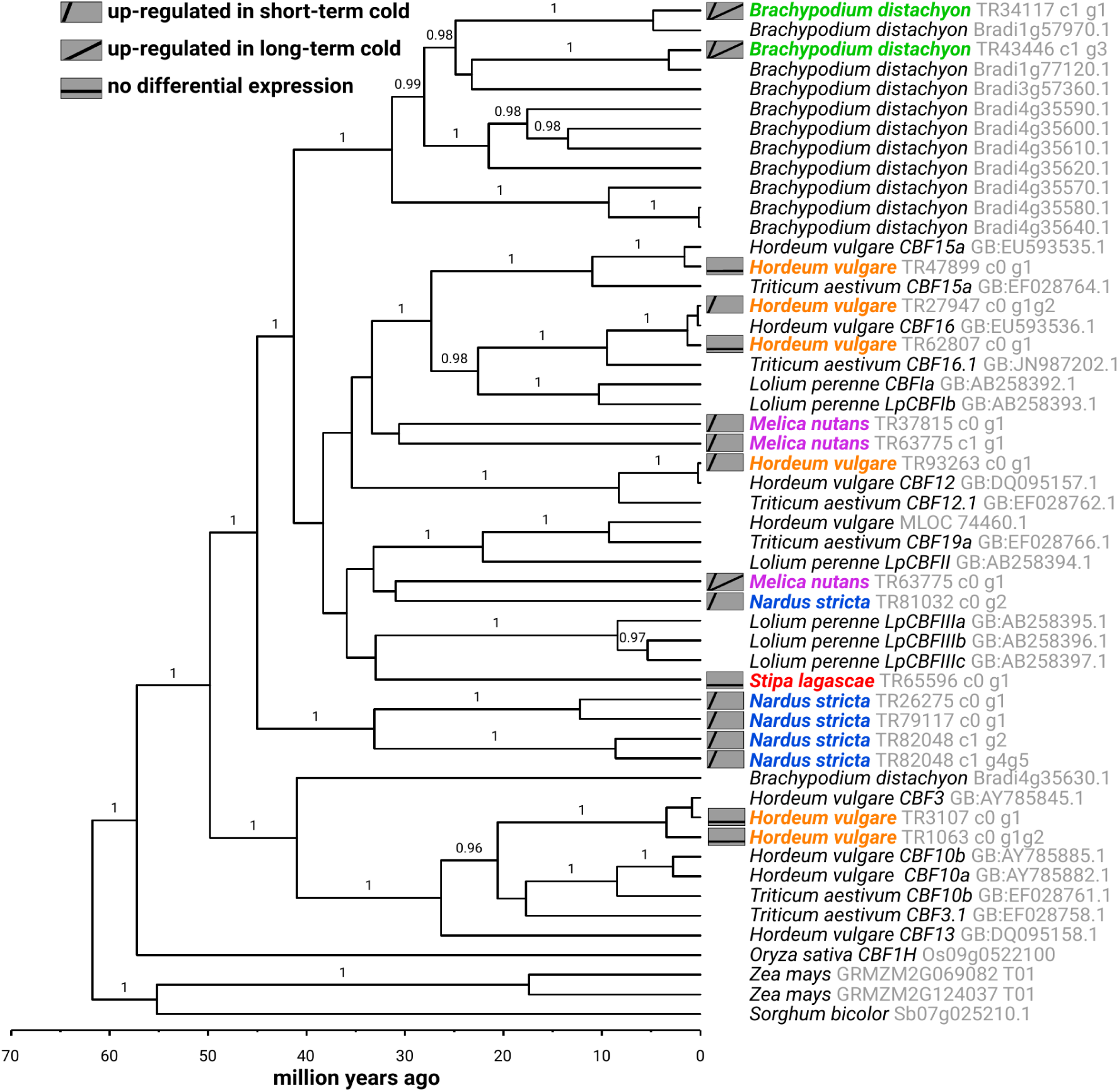
Time calibrated phylogeny for the Pooideae-specific CBFIIIc/d gene family. The phylogeny was estimated with BEAST v1.8.2 using a HKY+Γ and an uncorrelated, lognormal distributed molecular clock model. Significant posterior probabilities are shown as branch labels. CBFIIIc and CBFIIId form to distinct clades.

The CBFIV gene tree (Fig. 3) reconstructed the four known *CBFIV* clades (Badawi et al. 2007) but no homologous transcripts from *N. stricta* or *B. distachyon* were identified. Two short-term cold induced transcripts from *S. lagascae* and *M. nutans* formed a monophyletic clade together with the core Pooideae *CBFIV* homologs. In accordance with the fact that there are no known *CBFIVb* homologs in *H. vulgare*, we only identified cold induced *H. vulgare* transcripts in the three remaining *CBFIV* clades. Interestingly the two *de novo* transcripts from *M. nutans* and *S. lagascae* were induced by short-term cold.

**Figure 3:**
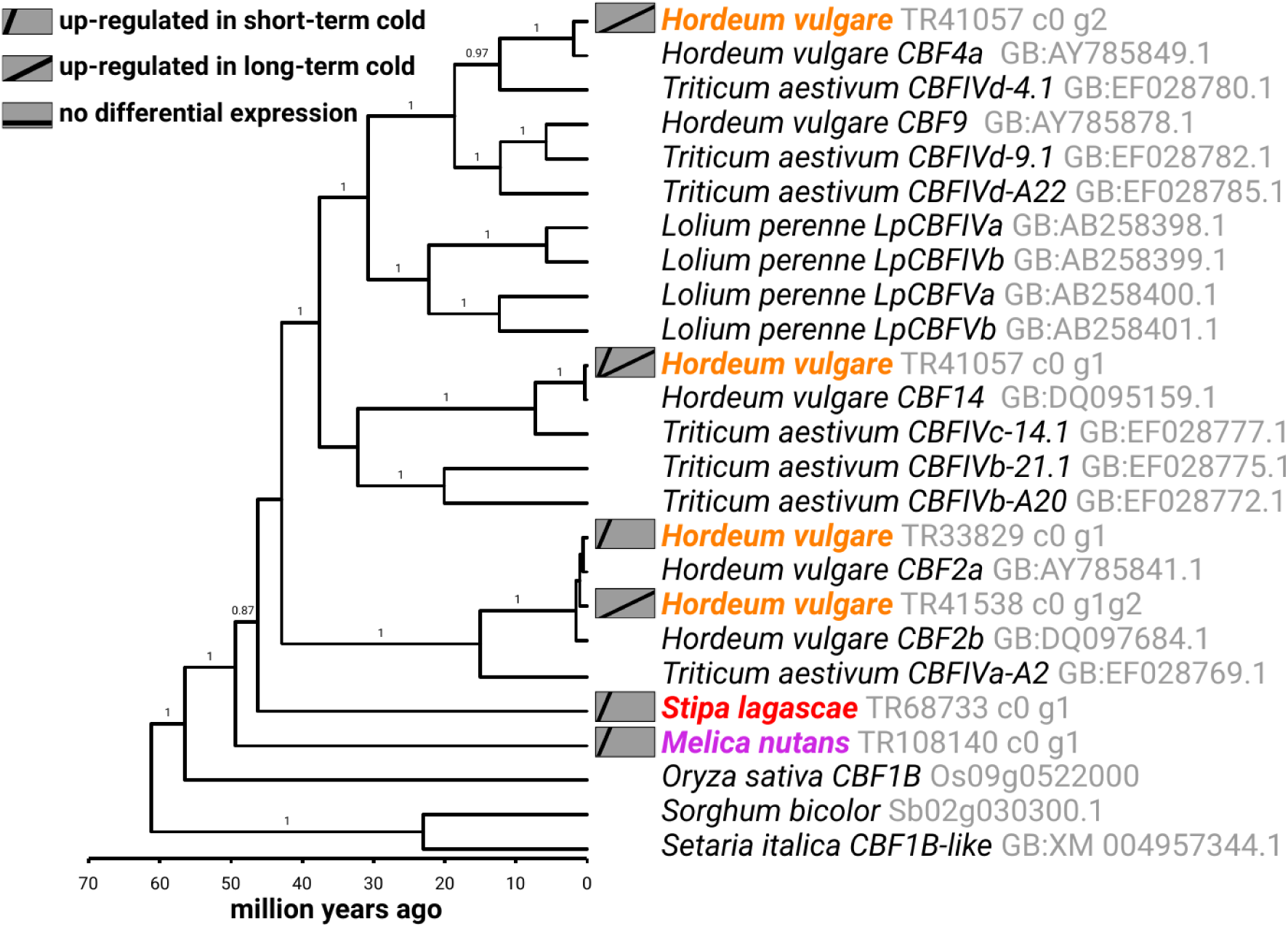
Time calibrated phylogeny for the Pooideae CBFIV gene family. The phylogeny was estimated with BEAST v1.8.2 using a HKY+Γ and an uncorrelated, lognormal distributed molecular clock model. Significant posterior probabilities are shown as branch labels.

### Dehydrin genes

Dehydrin (*DHN*) genes are well studied in *H. vulgare* and to date there are 13 known dehydrin homologs. Structurally they can be grouped into four distinct types based on the presence of amino acid segments (Y, K and S): SK_3_-type (Hv*DHN8*), KS-type (Hv*DHN13*), K_n_-type (Hv*DHN5*) and Y_n_SK_n_-type dehydrins (the 10 remaining Hv*DHN*-genes) (Kosová et al. 2007). Because these four groups represent phylogenetically distinct clades (Karami et al. 2013), we reconstructed individual gene trees for each group.

Cold induced accumulation of the gene *HvDHN5* and its wheat ortholog *WCS120* is regarded as marker for frost tolerant plants in *H. vulgare and T. aestivum* (Kosová et al., 2007). Beside a short- and long-term cold induced *DHN5* transcript in *H. vulgare*, we did not identify any *HvDHN5* homologs in the other investigated species. That is in accordance with the fact that K_n_-type dehydrins have previously only been described in *Triticeae* species.

Short-term cold induced transcripts homologous to *HvDHN8* and *HvDHN13* were present in all investigated Pooideae species (Fig. S1 and S2). Altough the gene trees did not reflect the species phylogenies, the Pooideae homologs formed monophyletic clades. The literature contains no reports of *HvDHN8* and *HvDHN13* homologs in species of the *Poeae* tribe, e.g. *L. perenne*, and no homologous sequences have been deposited to GenBank (confirmed with blast searches). Contrary to this expectation, we identified one *L. perenne* homolog for each gene respectively.

The remaining barley dehydrin genes (*HvDHN1*, *HvDHN2*, *HvDHN3*, *HvDHN4*, *HvDHN6*, *HvDHN7*, *HvDHN9*, *HvDHN10*, *HvDHN11* and *HvDHN12*) belong to the Y_n_SK_n_-type. All homologous *de novo* transcripts belonged to four clades (*HvDHN1-2, HvDHN9-12, HvDHN3-4-7* and *DHN10)*, which were specific to Pooideae (Fig. 4) and formed a clade with homologs from *O. sativa, Z. mays* and *S. bicolor*. Three of the Pooideae-specific clades contained transcripts from early diverging Pooideae species, with *DHN1-2* being specific to the core group and *B. distachyon*. While all transcripts of *N. stricta* and *M. nutans* were induced by short- and long-term cold treatment, *DHN* transcripts of *H. vulgare, B. distachyon* and *S. lagascae* were not differentially expressed.

**Figure 4:**
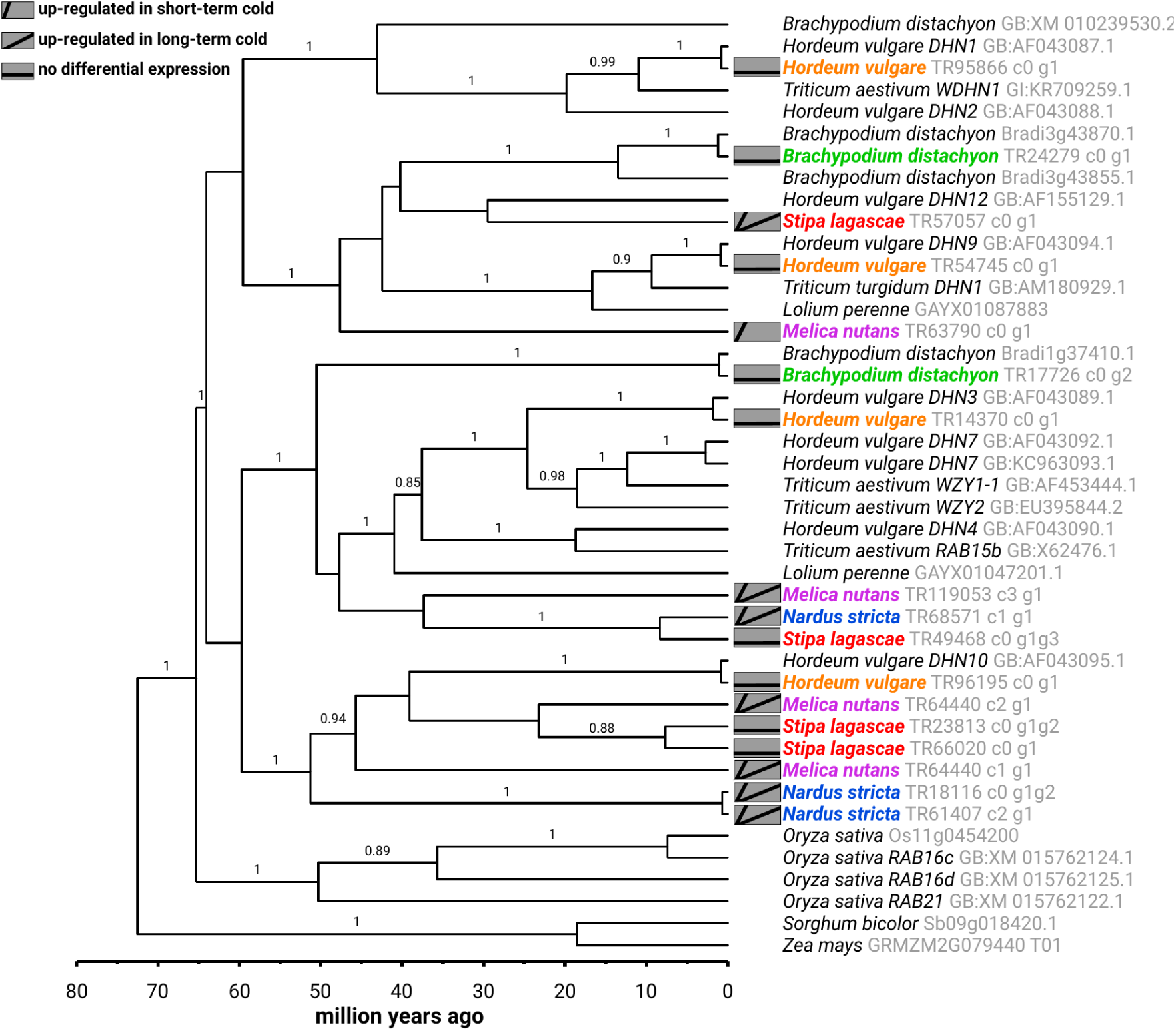
Time calibrated phylogeny for the Pooideae Y_n_SK_n_-type DHN gene family. The phylogeny was estimated with BEAST v1.8.2 using a GTR+Γ and an uncorrelated, lognormal distributed molecular clock model. Significant posterior probabilities are shown as branch labels.

For the last two Y_n_SK_n_-type clades – containing *HvDHN6* and *HvDHN11* – we did not identify any homologous, cold-induced transcripts, which was expected given that *HvDHN6* and *HvDHN11* are known to be expressed in embryos and caryopses only (Choi and Close 2000; Tommasini et al. 2008). These two clades were clearly less related to the rest of Y_n_SK_n_-type *DHN* homologs, because they were nested outside the monophyletic, Poaceae-specific Y_n_SK_n_-type clade (data not shown).

### Chloroplast cold regulated genes

The phylogenetic analysis of the *ctCOR* gene family reconstructed a monophyletic origin for all Pooideae *COR14*/*WCS19* homologs. Within this Pooideae-specific clade, two previously reported (Crosatti et al. 2013) sister clades contained core Pooideae homologs of *WCS19* and *COR14* (Fig. 5). However, their sister relationship was not well supported by the BEAST analysis. Those two clades were nested within a monophyletic clade, containing homologous transcripts from *B. distachyon, M. nutans* and *S. lagascae.* Except of one *S. lagascae* transcript, all other transcripts in this clade were induced by short- and long-term cold. The remaining homologs formed an unsupported clade, which contained one *N. stricta* transcript that was induced by short- and long-term cold.

**Figure 5:**
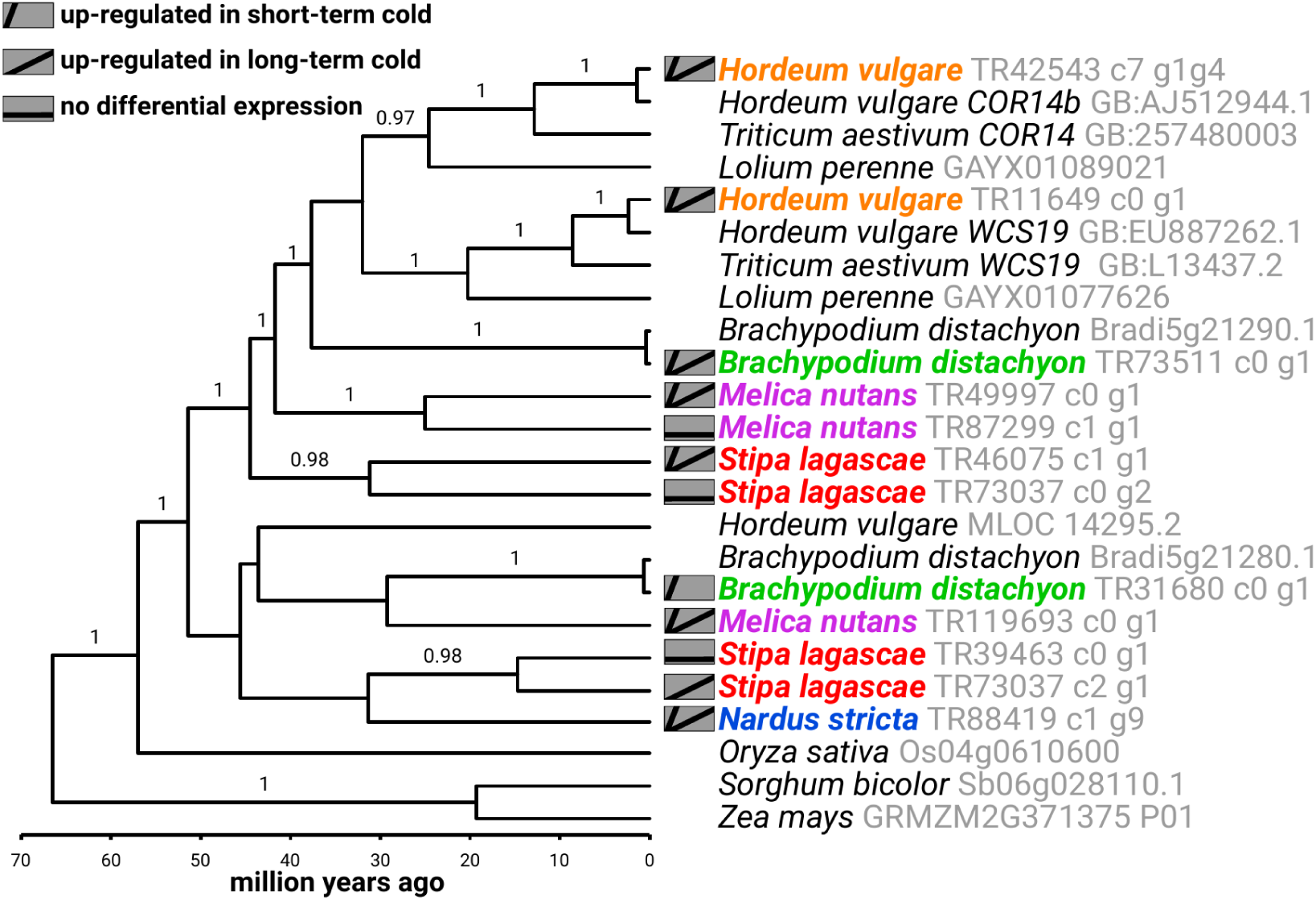
Time calibrated phylogeny for the Pooideae ctCOR gene family. The phylogeny was estimated with BEAST v1.8.2 using a GTR+Γ and an uncorrelated, lognormal distributed molecular clock model. Significant posterior probabilities are shown as branch labels.

### Ice-recrystallization inhibition protein genes

We identified six *H. vulgare,* three *B. distachyon* and one *S. lagascae* transcripts that formed a monophyletic clade (Fig. S3) with previously identified Pooideae *IRIP* homologs (Li et al. 2012). The *S. lagascae* transcript and one *H. vulgare* transcript were truncated and did not contain a characteristic IRI motif (see Li et al. 2012), even though the transcript from *S. lagascae* was induced by short- and long-term cold. The remaining transcripts conatained IRI motifs and were but one induced by short- and long-term cold. No IRIP homologs were identified for *M. nutans* and *N. stricta*.

### Fructosyltransferase genes

Phylogenetic analyses of the *FST* gene family reconstructed a monophyletic origin of the Pooideae *FST* homologs that contained two well supported main clades (Fig. S4). In the clade containing all known *FST* genes (Huynh et al. 2012) we identified two *H. vulgare* transcripts that were induced by long-term cold and contained the diagnostic FST motif (see Lasseur et al. 2008). Also part of that putative *FST* clade were one *H. vulgare,* one *B. distachyon* and one *M. nutans* transcript that lacked the FST motif. The other main clade contained putative vacuolar invertase 3 homologs of *H. vulgare, B. distachyon* and *N. stricta*.

## Discussion

### Cold acclimation evolved independently in different Pooideae lineages

Cold acclimation as adaptation to temperate climates is not restricted to core Pooideae species and *B. distachyon.* Our results (Fig. 1) show that cold acclimation is part of the cold adaptation in early-diverging Pooideae lineages. Cold acclimated plants of all tested species showed increased frost tolerance relative to non-acclimated plants. Interestingly, non-acclimated plants of the three early diverging Pooideae species exhibited high frost tolerance at 4°C relative to most tested core Pooideae and *Brachypodium* species. This might indicate that species of early diverging Pooideae possess a high tolerance against sudden frost shocks. This hypothesis however remains to be tested.

In core Pooideae, the gene *DHN5*, and genes from the families *FST*, *IRIP*, *CBFIV* and *CBFIII* are known to be crucially involved in cold acclimation and induced during short- and long-term cold (Choi et al. 2002; Vágújfalvi et al. 2003; Badawi et al. 2007; Knox et al. 2008; Zhang et al. 2010; Livingston et al. 2009; Knox et al. 2010; Soltesz et al. 2013; Jeknić et al. 2014; Todorovska et al. 2014; Marozsán-Tóth et al. 2015). The expression patterns of respective homologous *H. vulgare* transcripts presented here are in line with this research. However, the lack of conserved expression patterns of these genes and gene families among the tested Pooideae species indicates that cold acclimation is regulated differently in the five Pooideae lineages. These results contradict the hypothesis that key cold acclimation genes were recruited early in the Pooideae lineage. We therefore propose that regulation of cold acclimation evolved independently in Pooideae lineages and different genes were recruited.

For the *H. vulgare* cold acclimation gene *DHN5* and its ortholog in *T. aestivum WCS120*, no homologous transcripts were identified outside the Triticeae tribe and are neither reported in the literature (Kosová et al. 2012). Both genes belong to the K_n_-type dehydrins, that are characterized by multiple copies of the dehydrin specific K-segment. The lack of any K_n_-type dehydrins in other Pooideae species suggest that this specific family evolved relatively recently in the Triticeae – likely by expanding the K-segments of a dehydrin-like gene – and gained a crucial role in cold acclimation of Triticeae species (reviewed in Kosová et al. 2007, Kosová et al. 2012).

Two of the gene families whose function in cold acclimation is well described for core Pooideae are the *FST* and *IRIP* gene families (reviewed by Sandve et al. 2011), but no homologs were expressed in species of early-diverging lineages (Fig. 6). Although we identified *FST* homologs in *M. nutans* and *B. distachyon* (Fig. S4) and an *IRIP* homolog in *S. lagascae* (Fig. S3), the lack of diagnostic FST or IRI motifs suggests that functional genes are restricted to core Pooideae and core Pooideae-Brachypodieae, respectively. Hence, both gene families were recruited into cold acclimation in those two clades and represent more recent cold adaptations.

**Figure 6:**
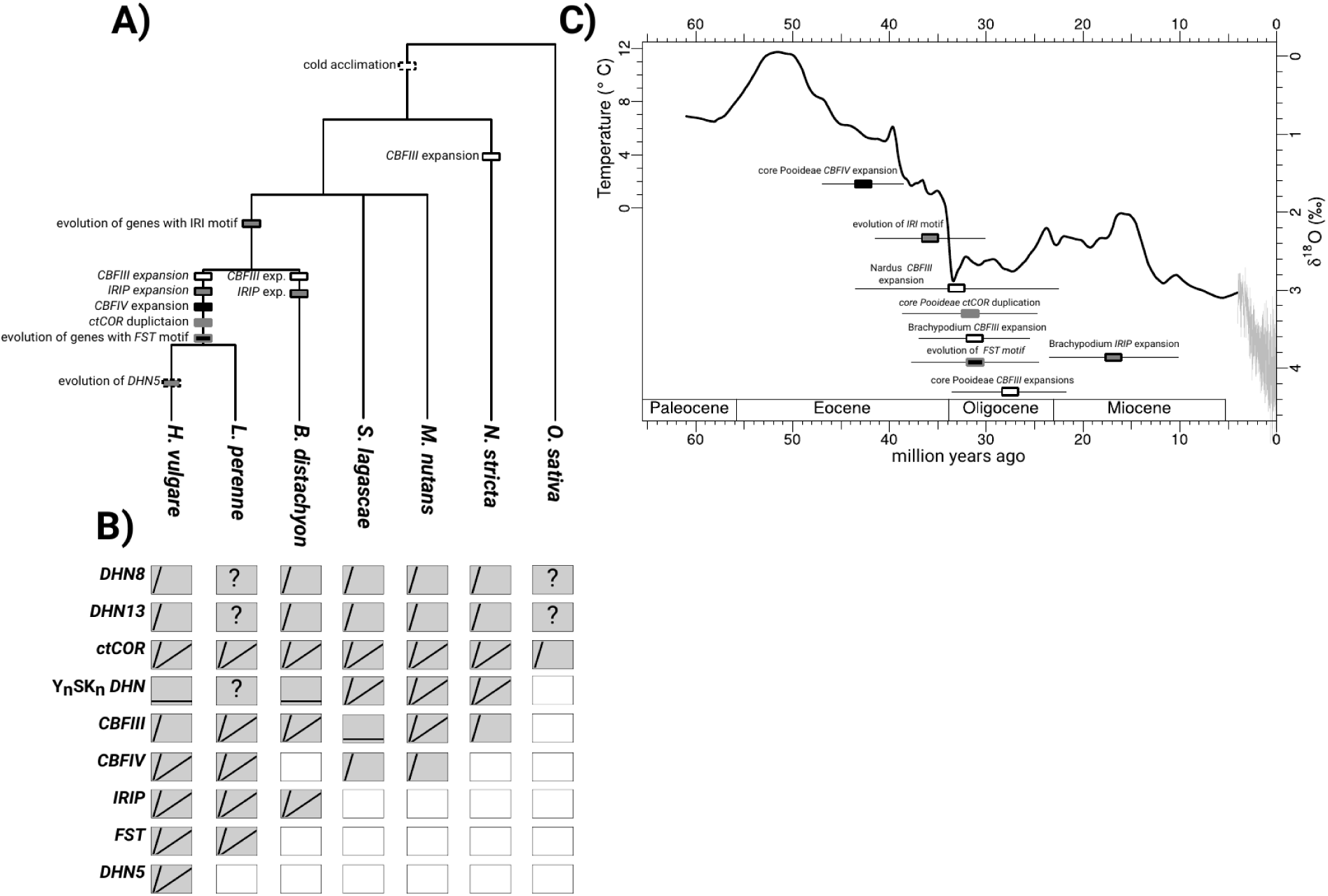
Summary of bhylogenic analyses, gene expression experiments and timeline of lineage-specific expansions. **A)** Emergence and expansion events of cold responsive gene families in relation to schematic Pooideae species tree. **B)** Long- and short-term cold responsive expression profiles of *de novo* transcripts for each of the focal species depicted above. Quesstion marks indicate that the expression pattern of the respective transcript is unknown, empty white boxes indicate that homologous transcripts were not identified or are not known. Expression patterns are approximated from the literature for *L. perenne* (*CBF* [Tamura and Yamada 2006], *WCS19/COR14* [Oishi et al. 2010], *FST* [Hisano et al. 2008; Paina et al. 2014]) and O. sativa (*ctCOR-like* [Maruyama et al. 2014]). **C)** Inferred time period for emergence and expansion events depicted in A). δ^18^O (%o) values for times without sea ice, which roughly applies to the period before the temperature shown on left y-axis applies for times without sea ice, which roughly applies to the Eocene-Oligocene transition. Whiskers represent 95% of the highest posterior density (HPD) distribution.

Genes of the *CBFIII* and *CBFIV* families code for transcription factors that are important for regulation of cold acclimation in core Pooideae and in case for *CBFIII* for *B. distachyon* (Badawi et al. 2007, Li et al. 2012). Our analyses could not resolve the species topology within the *CBFIII* gene tree (Fig. 2), but we identified homologous transcripts in all early diverging Pooideae species that were cold induced. Except for one *M. nutans* transcript however, we suggest that *CBFIII* homologs are not involved in cold acclimation of early diverging Pooideae species, because they are not induced by long-term cold treatment. Homologs closely related to *CBFIV* have never been reported for *B. distachyon* and also our results lack evidence for such transcripts. Although we identified *CBFIV* transcripts (Fig. 3) in *S. lagascae* and *M. nutans,* they were not induced by long-term cold and are unlikely to be involved in cold acclimation. We therefore suggest that *CBFIV* genes were recruited into cold acclimation in core Pooideae only, correlating with the expansion of this gene family.

While Y_n_SK_n_-type *DHN* homologs are known to be expressed during frost in core Pooideae – which is in line with our findings for *H. vulgare* – transcripts from *M. nutans, S. lagascae* and *N. stricta* are induced by long-term cold (Fig. 4). These results suggest that those transcripts might be involved in cold acclimation exclusively in early-diverging Pooideae lineages. Functional studies are necessary to investigate their role in cold acclimation.

The only tested genes whose function in cold acclimation might be conserved throughout the Pooideae are homologs of the *ctCOR* gene family (Fig. 5), because transcripts in all investigated species were induced by long-term. The gene tree failed to reconstruct the species topology of the Pooideae, due to an unsupported clade containing the one *N. stricta* transcript. Two duplication events – one after the divergence of *N. stricta* and one in the core Pooideae giving rise to the *COR14* and *WCS120* clade – could explain the abundance and relationship of *ctCOR* homologs in a parsimonious way. The conserved expression pattern of the *ctCOR* homologs renders this gene family a very interesting candidate for further studies of the regulation of cold acclimation in Pooideae. Our results suggest that *ctCOR* genes might have been involved in a putative cold acclimation of the Pooideae MRCA.

Opposed to cold acclimation genes which are induced by long-term cold. Genes involved in response to short-term cold seem to be more conserved. Among dehydrin genes homologs of *HvDHN8* (Fig. S1) and *HvDHN13* (Fig. S2) exhibited the same expression pattern in all species, in addition to highly conserved nucleotide sequences and a lack of duplication events. Because all other gene families showed signs of lineage specific evolution, the findings for *DHN8* and *DHN13* suggest that their cold-responsive function has been conserved since the MRCA.

### Pooideae lineages evolved specific cold adaptation by expanding gene families

Using transcriptomic data to reconstruct phylogenetic relationships between genes is inherently problematic. Homologous genes might exist in early diverging Pooideae species, but might not be expressed. In addition, coding sequences might not contain sufficient informative substitutions to reconstruct the true species topology. Nonetheless we were able to identify homologous transcripts and argue that a lack of long-term cold induced transcripts correlates with lack of function in cold acclimation.

Furthermore the gene trees displayed several duplication events and gene family expansions. Based on our results we propose that gene family expansion was an important mode of cold adaptation in the Pooideae subfamily. Our results corroborate findings from Sandve and Fjellheim (2010), who identified an increase of gene copy number in *CBFIV*, *FST* and *IRIP* gene families as an evolutionary force of cold climate adaptation of core Pooideae and *Brachypodium* species. Although we lack sufficient genomic data for early diverging Pooideae species, we found evidence for Pooideae specific expansions in *CBFIII, Y_n_SK_n_-type DHN* and *ctCOR* gene families.

*Brachypodium distachyon* possesses long-term cold induced transcripts for the *IRIP* (Fig. S2) and also *CBFIIId* (Fig. 2) gene families. Both gene families expanded specifically in the *B. distachyon* lineage and we therefore argue that their function in cold acclimation evolved independently from the core Pooideae. The Y_n_SK_n_-type *DHN* family (Fig. 4) expanded into at least three distinct clades early in the Pooideae history, possibly in the MRCA. Another Y_n_SK_n_-type *DHN* clade (containing *HvDHN1* and *2*) seems to have emerged in the Brachypodieae and core Pooideae. Due to an unresolved gene tree it is difficult to assess where in the Pooideae phylogeny *CBFIII* gene family began to expand (Fig. 2), but it is apparent that there were at least three independent expansions in the Nardeae tribe, Brachypodieae tribe and the core Pooideae that lead to several gene copies. Expansion of gene families may have lead to functional specialization or novel functions of the various gene copies. Other studies have shown that stress related gene families tend to expand via tandem duplications (Hanada et al. 2008), which may lead to lineage-specific expansion of the gene family (Lespinet et al. 2002).

Our estimates placed the evolution of FST and IRI motifs as well as the core Pooideae specific duplication of the *ctCOR* family and *CBFIII* expansions in the period of the E-O transition (Fig. 6). Most interestingly, two independent expansions of *CBFIII* the lineage of *N. stricta* and *B. distachyon* happened during the same time period. Those findings partly confirm analyses by Sandve and Fjellheim (2010), who suggested that cold responsive gene families expanded due to increased selection pressure for improved cold adaptation during the E-O transition. In contrast to Sandve and Fjellheim (2010), we were unable to correlate initial core Pooideae-specific expansions of the *CBFIV* family with the E-O transition. In the case of *FST* and *IRIP* gene families the increased cold stress during the E-O transition might have led to the evolution of novel protein domains in the core Pooideae and Brachypodieae (Sandve et al. 2008; Li et al. 2012). It remains to be tested if similar novelties evolved in other lineages. Based on our findings, there is evidence that the E-O transition affected the molecular evolution of cold adaptive mechanisms in all investigated Pooideae lineages. By the time of the E-O transition, all major Pooideae lineages had already diverged (Marcussen et al. 2014; Grønvold et al. 2017) and this supports the indications in our data that cold adaptation largely evolved separately in different Pooideae lineages.

### Drought tolerance – an evolutionary basis for cold tolerance?

Parts of the early cold tolerance in Pooideae might have been derived from ancestral drought tolerance. Cold stress may cause dehydration due to decreased membrane stability as well as lead to accumulation of ROS (Kratsch and Wise 2000; Murata et al. 2007; Crosatti 2013), similar to stresses occurring during drought (Mahajan and Tutej 2005). Following this, several authors have speculated about the importance of drought tolerance for the early evolution of the Pooideae (Kellogg 2001; Schardl et al. 2008; Vigeland et al. 2013).

Due to their molecular functions in membrane stabilizing and ROS scavenging, ancestrally drought-responsive genes were suitable candidates for cold responsive pathways when ancestral Pooideae species faced temperate conditions. It has been shown that genes with suitable, pre-existing molecular functions seem to be recruited into certain molecular pathways preferentially (True and Carroll 2003; Christin, Osborne et al. 2013; Christin et al. 2015). Some of our findings lend support to this scenario. Firstly, early-diverging lineages possess cold-induced transcripts of *CBFIII* (Fig. 2) and *CBFIV* (Fig. 3) and many *CBF* genes are known to be involved in drought tolerance in several angiosperms (Agarwal et al. 2006; Akhtar et al. 2012). Secondly, *ctCOR* (Fig. 5) transcripts are cold induced in all species. The *ctCOR* genes *COR14* and *WCS19* are thought to be involved ROS-mediated stress (Crosatti et al. 2013), which is both beneficial during drought and cold stress. Thirdly, *DHN8* (Fig. S1) is an ancestral drought-responsive gene, that gained a function in cold tolerance. Orthologs of *DHN8* are involved in the protection of plasma membrane during cold (Yang et al. 2014) and drought stress (Danyluk et al. 1998; Houde et al. 2004), due to a putative ROS scavenging function (Kumar et al. 2014). Since its drought responsiveness seems to be conserved outside the Pooideae (Lee et al. 2005; Badicean et al. 2012), it is likely that *DHN8* gained its cold responsiveness first in the Pooideae MRCA.

## Conclusion

Taken together, our results provide valuable insights in the early adaptive evolution of Pooideae and contribute to the understanding how cold acclimation evolves in plants. We found signs of conserved cold response that likely existed in the Pooideae MRCA and increased the Pooideae’s potential to evolve cold tolerance. The conserved fraction of cold response might have enabled early Pooideae members to survive the first encounters with temperate conditions, making subsequent cold adaptation possible. Even though all species were able to increase their frost tolerance in response to cold acclimation, we observed a trend of lineage specific evolution of the regulation of cold acclimation. This has led to increased gene family complexity by gene family expansion, particularly within the core Pooideae lineage (Fig. 6).

Due to the scarce fossil record, divergence times of early-diverging Pooideae lineages are still under dispute and so is the stem age of the Pooideae. In the absence of reliable fossil calibration points our results contribute to a better understanding of climatic changes, that influenced molecular evolution of certain gene families in Pooideae subfamily. However, more reliable calibration points and a resolved genome-level phylogeny of the Pooideae subfamily will help us to improve the dating of the evolutionary history of cold adaptation. This will enable us to confidently correlate paleoclimatic events, like the E-O transition, with molecular innovations in order to reconstruct the colonization of temperate biomes by the Pooideae subfamily. And lastly, higher resolution, i.e. higher phylogenetic coverage, of cold adaptation evolution in the core Pooideae lineage will be valuable to understand the molecular traits that might have contributed to their putative rapid radiation, species richness and expansion into extreme habitats.

## Acknowledgments

The presented research was funded by the Nansen Foundation and through the TVERRforsk grant provided to SF, TRH and SRS by the Norwegian University of Life Sciences (NMBU). This work was part of the PhD project of MS and LG funded by NMBU. We thank Åsmund Bjørnstad, Morten Lillemo, Mallikarjuna Rao Kovi and USDA-NPGS for providing seeds of *H. vulgare*, *T. aestivum*, *L. perenne* and *S. lagascae*, respectively. For technical assistance handling plants during growth experiments we thank Øyvind Jørgensen. We are grateful to Pascal-Antoine Christin for invaluable advice and to Erica Leder, Thomas Marcussen, Ursula Brandes, Camilla Lorange Lindberg and Martin Paliocha for helpful comments on earlier versions of this manuscript.

## Supplementary figures

**Figure S1:**
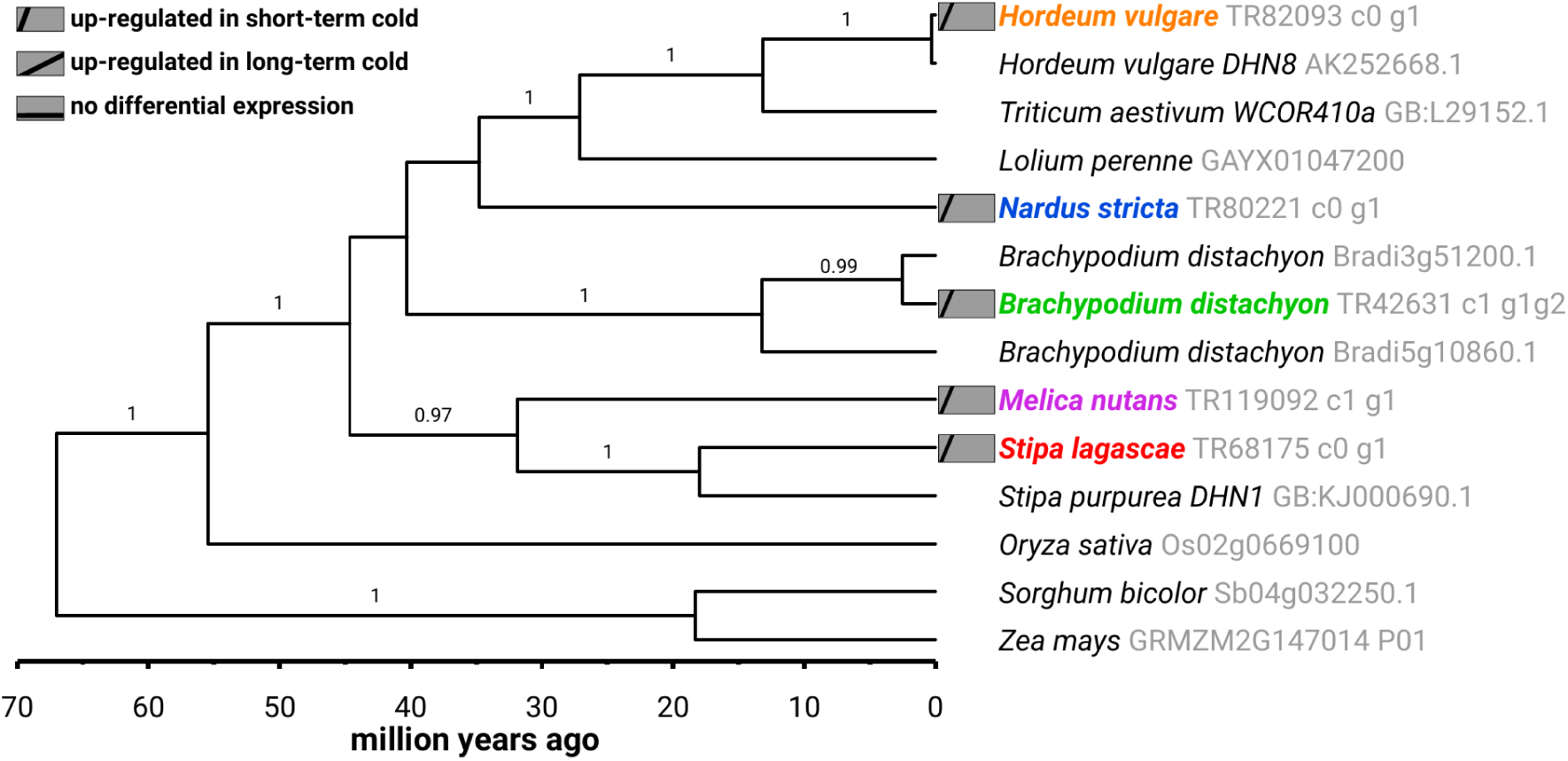
Time calibrated phylogeny for the Pooideae DHN8 gene family. The phylogeny was estimated with BEAST v1.8.2 using a GTR+Γ and an uncorrelated, lognormal distributed molecular clock model. Significant posterior probabilities are shown as branch labels.

**Figure S2:**
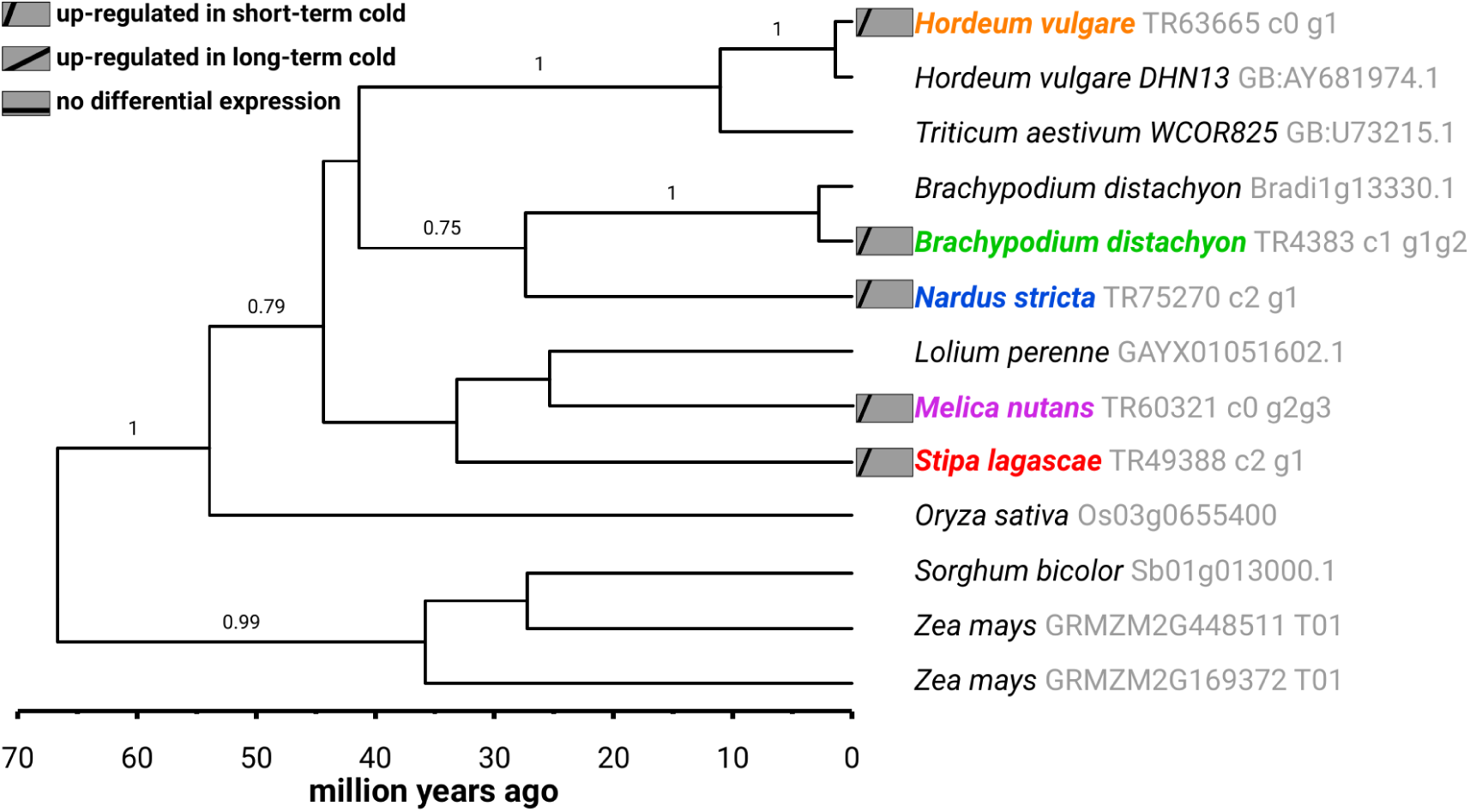
Time calibrated phylogeny for the Pooideae DHN13 gene family. The phylogeny was estimated with BEAST v1.8.2 using a GTR+Γ and an uncorrelated, lognormal distributed molecular clock model. Significant posterior probabilities are shown as branch labels.

**Figure S3:**
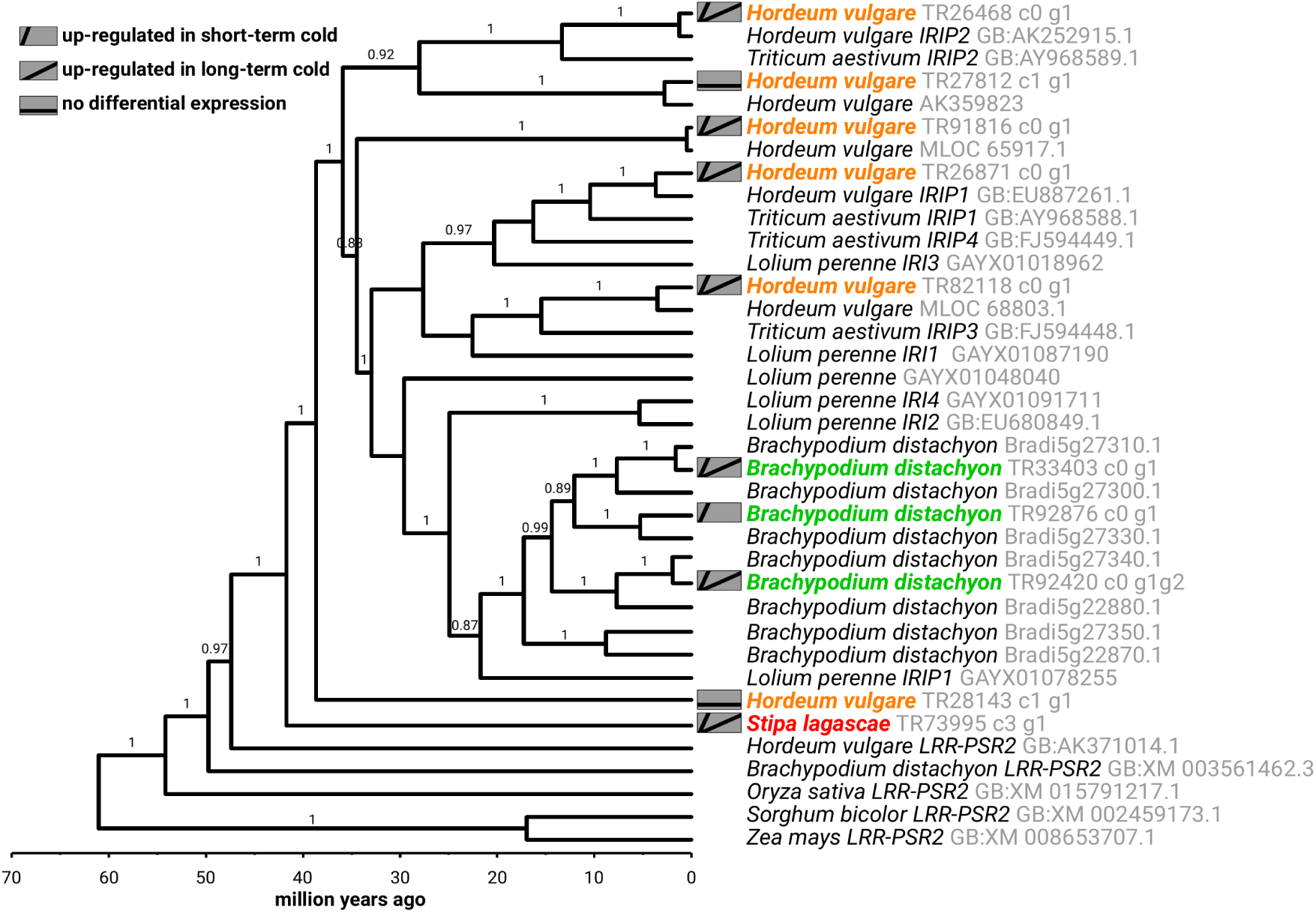
Time calibrated phylogeny for the Pooideae IRIP gene family. The phylogeny was estimated with BEAST v1.8.2 using a GTR+Γ and an uncorrelated, lognormal distributed molecular clock model. Significant posterior probabilities are shown as branch labels.

**Figure S4:**
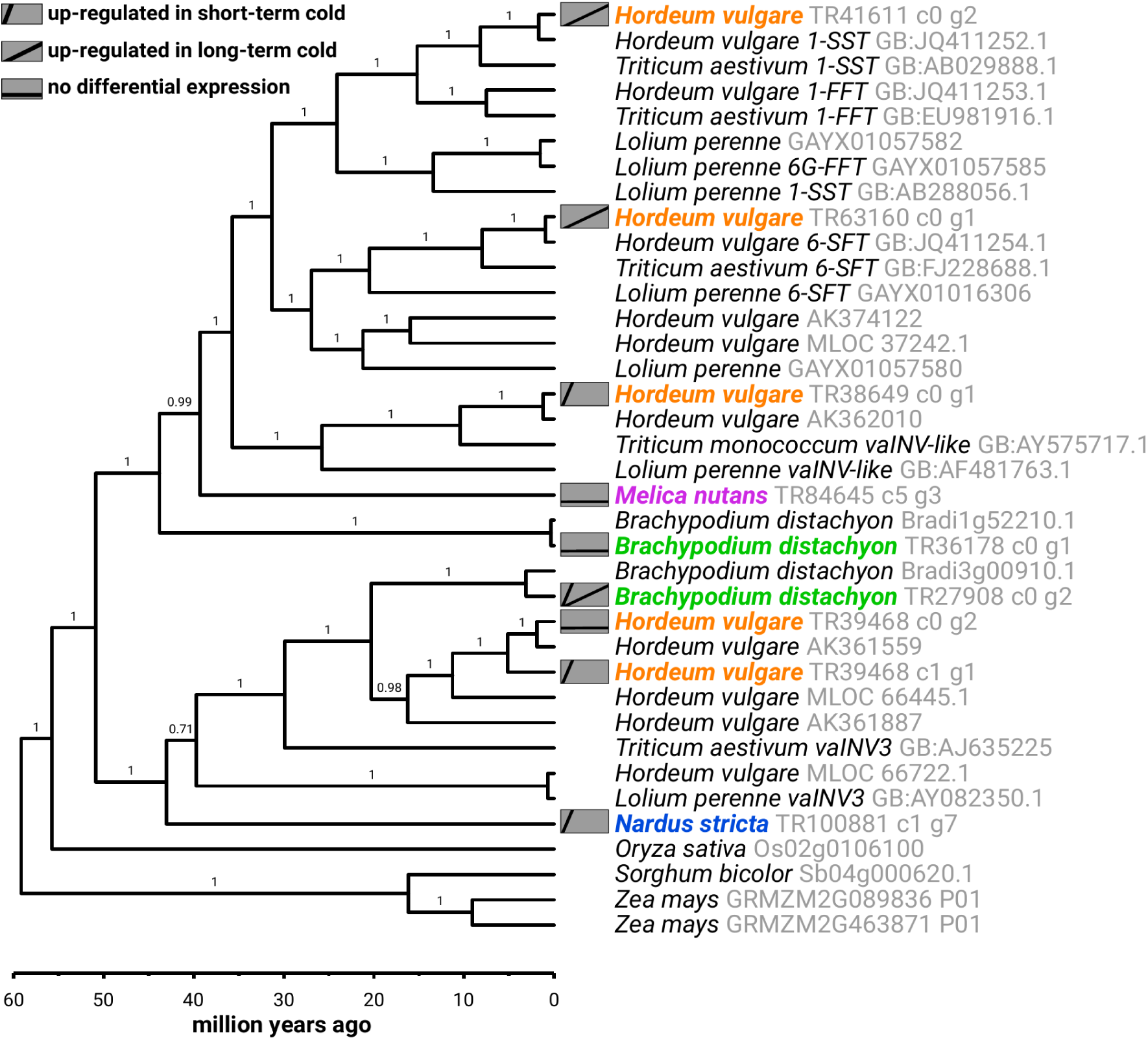
Time calibrated phylogeny for the Pooideae FST gene family. The phylogeny was estimated with BEAST v1.8.2 using a GTR+Γ and an uncorrelated, lognormal distributed molecular clock model. Significant posterior probabilities are shown as branch labels.

